# Tissue-specific dynamic codon redefinition in Drosophila

**DOI:** 10.1101/2020.06.22.165522

**Authors:** Andrew M. Hudson, Gary Loughran, Nicholas L. Szabo, Norma M. Wills, John F. Atkins, Lynn Cooley

## Abstract

Stop codon readthrough during translation occurs in many eukaryotes, including Drosophila, yeast, and humans. Recoding of UGA, UAG or UAA to specify an amino acid allows the ribosome to synthesize C-terminally extended proteins. We previously found evidence for tissue-specific regulation of stop codon readthrough in decoding the Drosophila *kelch* gene, whose first open reading frame (ORF1) encodes a subunit of a Cullin3-RING ubiquitin ligase. Here, we show that the efficiency of *kelch* readthrough varies markedly by tissue. Immunoblotting for Kelch ORF1 protein revealed high levels of the readthrough product in lysates of larval and adult central nervous system (CNS) tissue and larval imaginal discs. A sensitive reporter of *kelch* readthrough inserted after the second *kelch* open reading frame (ORF2) directly detected synthesis of Kelch readthrough product in these tissues. To analyze the role of cis-acting sequences in regulating *kelch* readthrough, we used cDNA reporters to measure readthrough in both transfected human cells and transgenic Drosophila. Results from a truncation series suggest that a predicted mRNA stem-loop 3’ of the ORF1 stop codon stimulates high-efficiency readthrough. Expression of cDNA reporters using cell type-specific Gal4 drivers revealed that CNS readthrough is restricted to neurons. Finally, we show that high-effficiency readthrough in the CNS is common in Drosophila, raising the possibility that the neuronal proteome includes many proteins with conserved C-terminal extensions. This work provides new evidence for a remarkable degree of tissue- and cell-specific dynamic stop codon redefinition in Drosophila.

## Introduction

Context-dependent word meaning has a genetic counterpart in the dynamic redefinition of stop codons in specific locations. Dynamic codon reassignment occurs when competition between two alternative outcomes during translational decoding produce two distinct products in proportions that reflect the balance of competition. Gene expression is enriched by dynamic stop codon redefinition since’ in addition to synthesis of protein encoded by the first open reading frame (ORF1), some of the product has a C-terminal extension. The extent of UGA, UAG or UAA readthrough depends on the identity of the stop codon and on its flanking sequence, especially the 3’ mRNA sequence. UGA and to a lesser extent UAG are most frequently used for productive readthrough. Instead of involving tRNA, standard decoding of UAG, UAA and UGA involves protein release factors and accessory proteins (1). A further distinctive feature is that in mammalian decoding, mammalian release factor 1 causes the nucleotide 3’ to the stop to be included in the ribosomal A-site (2, 3). Even where this feature does not occur, the identity of the base 3’ of a stop codon is relevant to its effectiveness for termination (4, 5).

Additional sequence, generally but not exclusively 3’, is relevant for the higher level of readthrough involved in evolutionarily-selected occurrences. Many functionally utilized readthrough cassettes in plant viruses have particular 6-9 nt sequences 3’ of the stop codon (6–10), and counterpart 3’ adjacent short sequences have been studied in detail in yeast (11) and mammals (6, 12–14). The proximity of these types of stimulators to the stop codon implies they mediate their effects while within the ribosomal mRNA entrance channel. Nearby 3’ intra-mRNA structures have long been known to be important for readthrough of the Drosophila gene, *headcase* (15), as has a pseudoknot for the readthrough of the Murine Leukemia Virus UAG gag terminator to yield the GagPol precursor that is the source of reverse transcriptase (16, 17). The effectiveness of the pseudoknot is aided by part of the readthrough product binding eRF1 and indirectly diminishing termination (18)).

Interestingly, regulation of readthrough of several plant viruses involves significant long-distance pairing of nucleotides just 3’ of the readthrough site with far distant complementary nucleotides, in several cases with considerable complexity (19–22). The far distant location of the 3’ component can have regulatory significance including avoidance of clashes between translating ribosomes and replicase (23, 24). Another study also showed the potential for relevant long distance pairing for several vertebrate alphaviruses (25).

On completion of genome sequencing of 12 divergent Drosophila species, bioinformatics analyses provided the first evidence for abundant readthrough in insects and one crustacean (26, 27). In a thorough study, Jungreis and colleagues estimated that the expression of more than 600 *A. gambiae* mosquito genes and 900 Drosophila genes involves functional readthrough (28). Insects and probably crustaceans may be the greatest users of functional readthrough and so be the counterpart of what certain ciliates, e.g. Euplotes, are for frameshifting (29) and what squids and octopuses are for mRNA editing (30). Ribosome profiling revealed additional instances of Drosophila readthrough and experimental data on readthrough efficiency (31). The reason for the expression of an unusually large number of insect genes utilizing readthrough is unknown. One issue posed by this is whether there are tissue-specific differences in the level of readthrough.

Despite the variety, and in several cases the elaborate nature, of stimulatory signals 3’ of readthrough stop codons, in general the efficiency of utilized readthrough is substantially less than that of programmed frameshifting counterparts. In 1993, a striking exception with very high levels of readthrough was reported in expression of the Drosophila *kelch* gene (32, 33). The Kelch protein produced by termination at the first stop codon is 689 aa. Like the first discovered occurrence of functional readthrough used by phage *Qβ* (34, 35), the *kelch* readthrough-derived C-terminal extension is long, 782 aa. The 689 aa Kelch protein is part of a ubiquitin ligase complex required for oocyte growth during oogenesis (36–38) and *kelch* mutants are female sterile (33). In addition to phenotypes during oogenesis, *kelch* mutations also result in defects in the larval peripheral nervous system, with loss of *kelch* causing a reduction in dendritic branching (39). Analysis of *kelch* expression revealed an unusually high level of *kelch* stop codon readthrough in imaginal discs (32) despite no known role for Kelch in that tissue.

Here we examine in detail where *kelch* stop codon readthrough occurs and investigate how mRNA structure contributes to readthrough efficiency. We report exquisite tissue specificity with readthrough occurring at elevated levels in the central nervous system (CNS) and imaginal discs. The full-length Kelch protein is abundant in larval and adult neurons, but not glial cells. We examined the contribution of cis-acting RNA features in stop codon readthrough efficiency and identified a predicted stem-loop as a readthrough enhancer. Finally, we provide evidence that efficient stop codon readthrough in the CNS is widespread in Drosophila.

## Results

### Tissue-specific stop codon readthrough of *kelch* in imaginal discs and CNS

Our previous analysis of *kelch* revealed developmental and tissue-specific regulation of stop codon readthrough, with a high level of readthrough observed in a sample containing both imaginal discs and associated larval CNS (32) We re-examined levels of *kelch* readthrough product in samples prepared from separated larval tissues and found the highest levels of readthrough product in both the CNS and imaginal discs with no detectable readthrough product in lysates of fat body, gut and salivary gland tissue (Fig. 1A). We examined *kelch* readthrough product in adult tissues and observed low levels in malpighian tubules, testis, and ovary in comparison with adult brain. Strikingly, adult brain samples exhibited high levels of *kelch* readthrough product, with approximately 70 of Kelch detected present as the readthrough product (Fig. 1A).

**Fig. 1.**
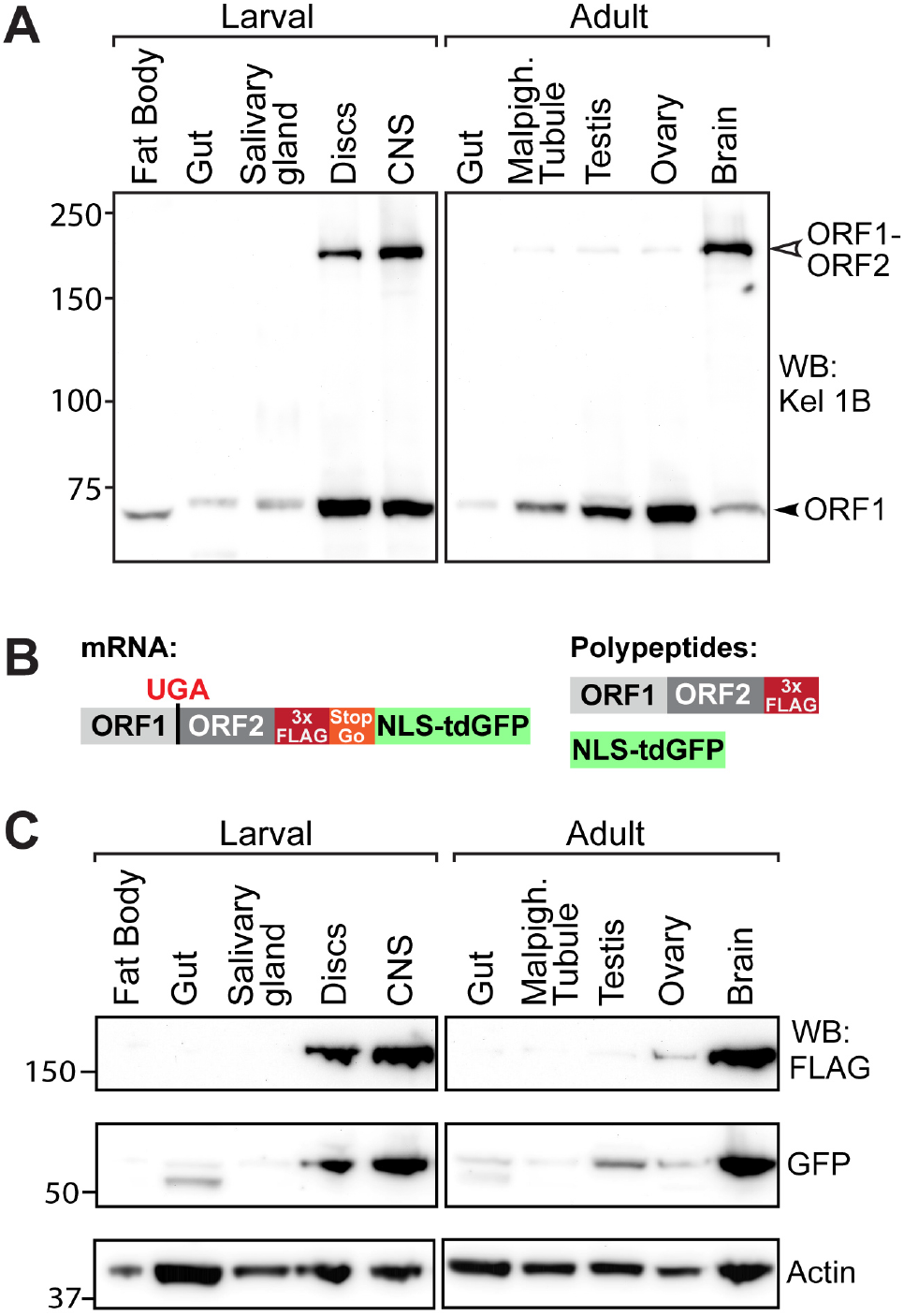
High-level readthrough of *kelch* mRNA in the Drosophila central nervous system. (A) Western blot of tissues dissected from 3rd-instar larvae and 3 – 5 day old adults probed with anti-Kelch. The predicted molecular masses for ORF1 and ORF1-ORF2 are 77 kDa and 160 kDa, respectively. (B) Schematic of reporter system for detecting readthrough of endogenous *kelch*. Translation through the ORF1 UGA stop codon results in the Kelch ORF1-ORF2 readthrough product tagged at its C-terminus with 3x FLAG and a distinct nuclear-localized tandem dimer GFP (NLS::tdGFP) polypeptide released during translation of a viral T2A/StopGo sequence. (C) Western blot of tissues dissected from 3rd-instar larvae and adult animals homozygous for the readthrough reporter insertion in the *kelch* locus, probed with *α*-FLAG to detect the readthrough product, *α*-GFP to detect the released NLS::tdGFP, and *α*-Actin as a loading control.

Our method to assess readthrough relied on an antibody against a Kelch antigen derived from the first open reading frame (ORF1) of *kelch* that detected both the ORF1 termination product and the readthrough product. Despite several attempts, we were unable to generate an antibody against antigens from the second open reading frame (ORF2) that would allow specific detection of the readthrough product. To investigate *kelch* readthrough more directly, we used CRISPR-Cas9-mediated homology-directed repair (HDR) to develop a reporter of endogenous *kelch* readthrough (Fig. 1B). This system allows for specific detection of the readthrough product using a 3xFLAG tag at the C-terminus of ORF2, and also results in the production of a discrete nuclear-localized tandem dimer GFP (NLS::tdGFP), released via a viral T2A/StopGo sequence (40, 41) following readthrough translation of ORF2. Importantly, the detection of the NLS::tdGFP protein served as a reliable assay of readthrough that is independent of the stability of the ORF1 or ORF1-ORF2 product. Western analysis of this reporter in adult and larval tissues showed elevated readthrough in the larval CNS and adult brain, with lower levels in the imaginal discs (Fig. 1C). A very low level of readthrough was observed in the ovary with both the *α*-GFP and *α*-FLAG antibodies, which is also seen in blots probed for Kelch (Fig. 1A). Interestingly, whereas GFP was present in the testis lysate, the FLAG antibody failed to detect ORF1-ORF2 protein in the testis sample, suggesting that the ORF1-ORF2 product is less stable in the testis than in other tissues.

To further characterize tissue-specific *kelch* readthrough expression, we imaged NLS::tdGFP produced by the readthrough reporter. For larval tissues, we compared the pattern of NLS::tdGFP expression in the CNS, where the readthrough product was highest by immunoblot, with salivary gland, which had no detectable readthrough product. Within the CNS, the highest levels of NLS::tdGFP were observed in cells of the central brain (CB), as well as additional cells in the ventral nerve cord (VNC) (Fig. 2A). Closer examination of the central brain revealed nuclear accumulation of NLS::tdGFP in clusters of cells adjacent to cells with no NLS::tdGFP (Fig. 2B – B’), suggesting cell-type specific regulation of *kelch* readthrough in the CNS. In contrast, no nuclear NLS::tdGFP was observed in salivary gland cells (Fig. 2C, D, D’), consistent with the lack of readthrough product observed in immunoblots.

**Fig. 2.**
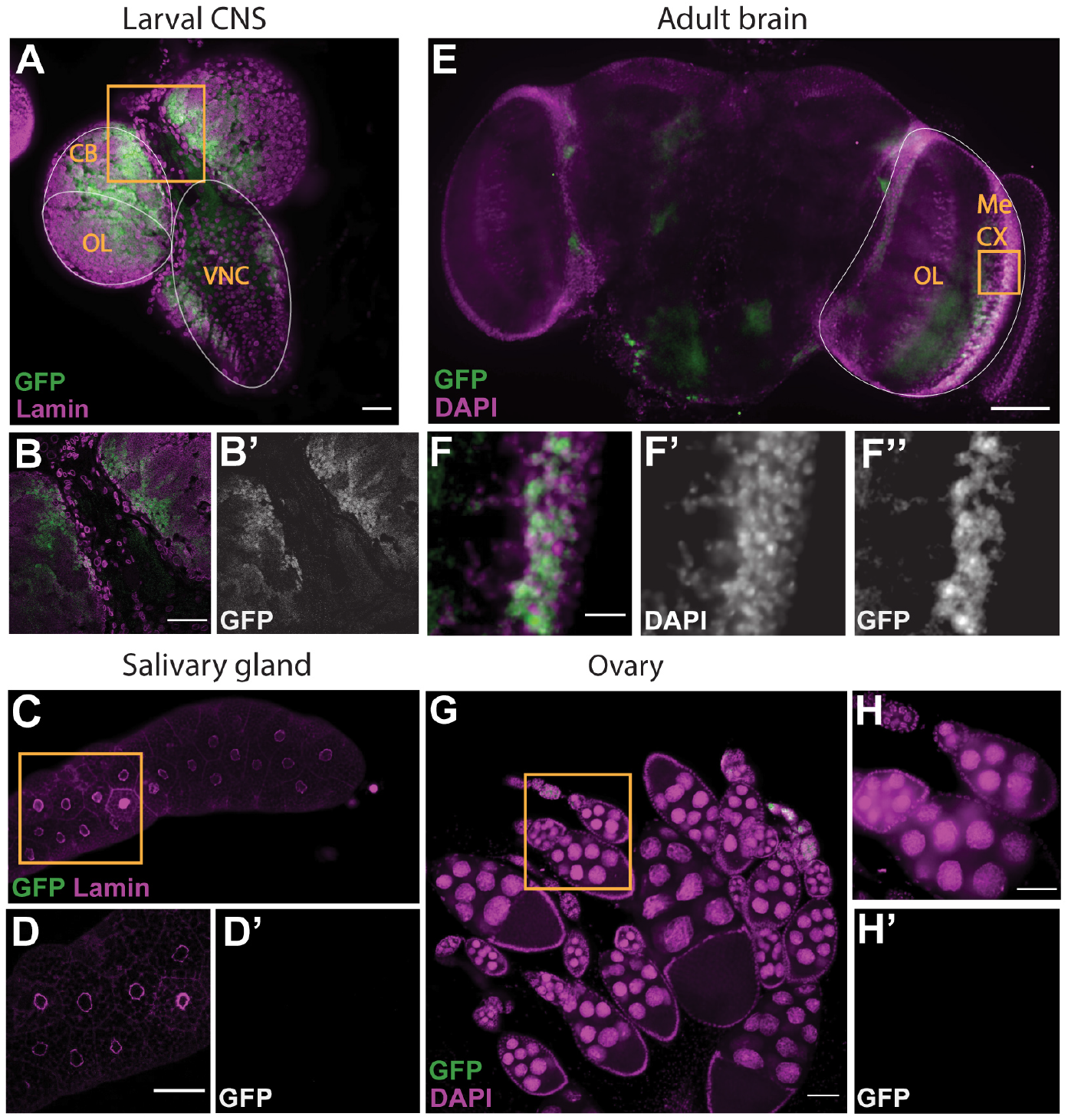
Visualization of endogenous *kelch* readthrough. Four tissues were analyzed for the presence of NLS::tdGFP produced following *kelch* readthrough translation: (A, B) larval CNS, (C, D) larval salivary gland, (E, F) adult brain, and (G, H) adult ovary. GFP levels in salivary glands (C-D) were normalized relative to those in larval CNS (A-B). Similarly, GFP levels in ovary (G-H) were normalized relative to the adult brain (E-F). Orange boxes in A, C, E, and G mark regions shown at higher magnification in lower panels. NLS::tdGFP accumulated in distinct subsets of cells of the larval (B – B’) and adult (F – F”) CNS, but not in salivary gland (D – D’) or ovary (H-H’). Scale bars for A-E and G-H = 50 *μm*. Scale bar for adult brain high magnification image (F – F”) =10 *μ*m. CB: central brain; OL: optic lobe; VNC: ventral nerve cord; Me CX: medulla cortex.

To further characterize tissue-specific *kelch* readthrough expression, we imaged NLS::tdGFP produced by the readthrough reporter. For larval tissues, we compared the pattern of NLS::tdGFP expression in the CNS, where the readthrough product was highest by immunoblot, with salivary gland, which had no detectable readthrough product. Within the CNS, the highest levels of NLS::tdGFP were observed in cells of the central brain (CB), as well as additional cells in the ventral nerve cord (VNC) (Fig. 2A). Closer examination of the central brain revealed nuclear accumulation of NLS::tdGFP in clusters of cells adjacent to cells with no NLS::tdGFP (Fig. 2B – B’), suggesting cell-type specific regulation of *kelch* readthrough in the CNS. In contrast, no nuclear NLS::tdGFP was observed in salivary gland cells (Fig. 2C, D, D’), consistent with the lack of readthrough product observed in immunoblots.

In adults, NLS::tdGFP was observed in cortical cells throughout the brain (Fig. 2E), including prominent labeling of cells within the medulla cortex (Me CX) in the optic lobe (boxed region in Fig. 2E). At higher magnification, comparison of NLS::tdGFP with DAPI revealed that NLS::tdGFP accumulated in only a subset of cells in the medulla cortex (Fig. 2F – F”). Examination of adult ovaries revealed no detectable NLS::tdGFP accumulation (Fig. 2G, H – H’), similar to larval salivary gland and consistent with the low-level readthrough product observed by immunoblot. Together, these results suggest a striking degree of tissue- and cell-specific regulation of *kelch* stop codon readthrough.

### Sequences 3’ of the UGA codon stimulate readthrough

There are several examples of stop codon readthrough stimulation by proximal RNA secondary structures (17, 19, 25, 43). In *kelch* mRNA, a predicted stem loop (SL1) starting eight nt 3’ of the ORF1 stop codon is supported by nucleotide conservation (Fig. 3C and Fig. S1; (25)). In addition, we identified the potential for a second stem loop (SL2) beginning 31 nt 3’ of the ORF1 stop (Fig. 3C) that is also widely conserved in Drosophila (Fig. S1).

**Fig. 3.**
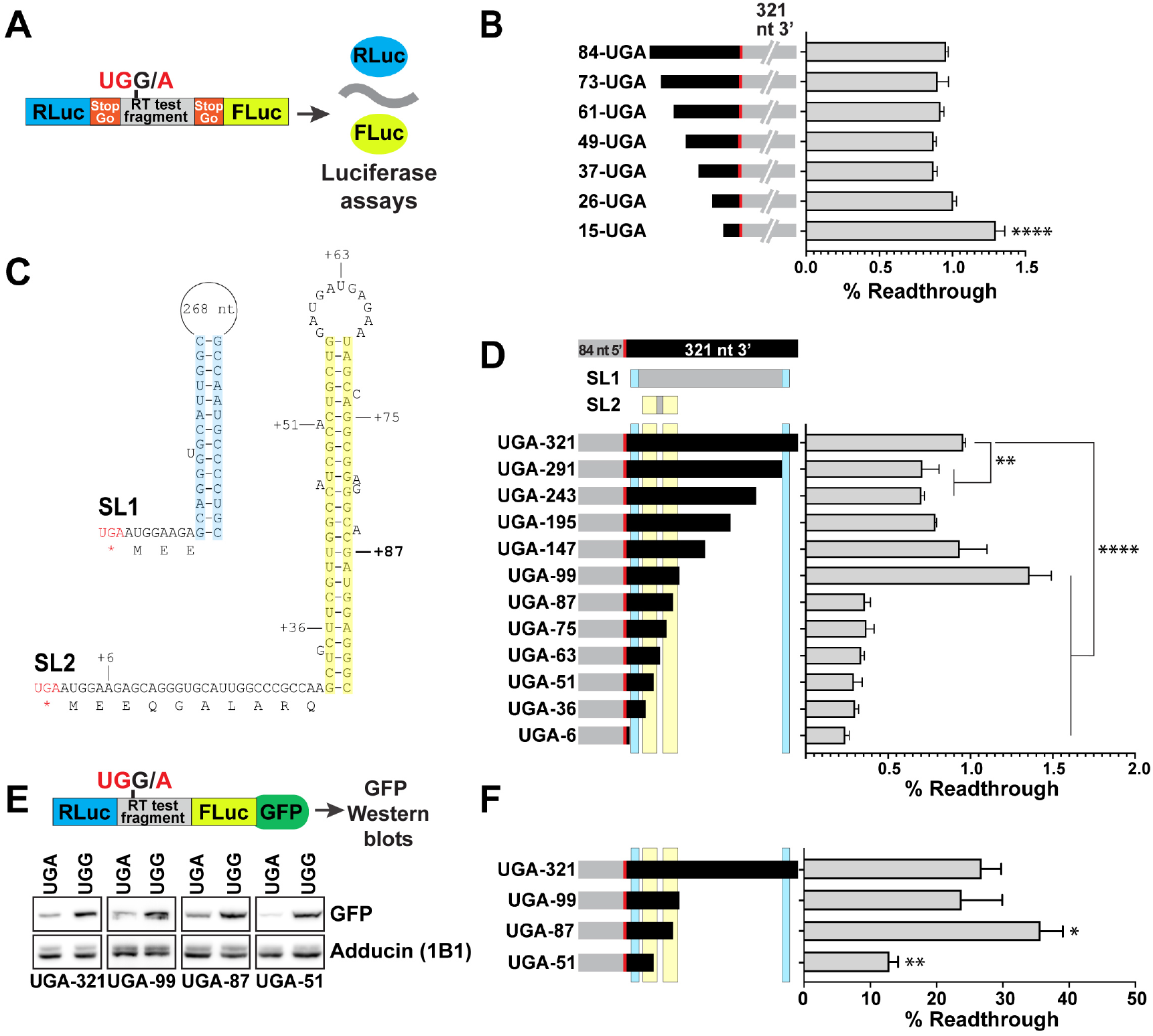
Analysis of the contribution to readthrough of *cis*-acting mRNA structural elements near the *kelch* stop codon. (A) Diagram of pSGDluc reporter construct. StopGo sequences from foot and mouth disease virus (FMDV) flanking the readthrough test fragment resulted in the translation of luciferase polypeptides free of residues encoded by *kelch* sequence; readthrough efficiency was calculated based on measured luciferase activities (42). (B) Sequences 5’ of the *kelch* stop codon were analyzed by transfecting the indicated 5’ truncation constructs into HEK-293T cells and measuring readthrough efficiency. (C) Diagram of predicted stem loop structures 3’ of the *kelch* ORF1 UGA codon. SL1 was previously described (25). For SL2, residues corresponding to the limits of 3’ deletions in (D) are indicated; location of 87 nt 3’ UTR is marked in bold. (D) Sequence requirements 3’ of the *kelch* UGA were analyzed as in (B) using the indicated 3’ deletion constructs. Predicted stem loops SL1 and SL2 are indicated above, with cyan or yellow shading corresponding to the shaded sequences in (C). In B and D, n = 3 biological replicates. (E) Diagram of dual luciferase construct for analysis of readthrough in transgenic Drosophila. GFP coding sequence was fused to the Renilla luciferase gene and readthrough efficiency was calculated from the ratio of GFP produced by UGA constructs compared to matched in-frame UGG controls. Constructs were expressed in adult brains using *nSybGal4* and lysates were prepared from heads. Representative western blots of constructs analyzed are shown below the diagram. (F) Quantification of readthrough efficiencies in flies; n ≥ 3. In B, D, and F, mean and SD are plotted. Readthrough efficiencies of truncation constructs were compared to the corresponding in-frame control using ANOVA with multiple comparisons correction. * indicates p < 0.05, ** p < 0.01, **** p < 0.0001.

To test the possible stimulatory role of *kelch* sequences both upstream and downstream of the ORF1 stop codon, we generated a series of dual luciferase reporters with 5’ and 3’ deletions and then assessed readthrough in HEK-293T cells (Fig. 3A-D). We observed a readthrough efficiency of 1% from *kelch* reporters with 84 nt 5’ and 321 nt 3’, which encompasses the predicted stem loop structures. We did not detect a decrease in readthrough efficiency with any of the 5’ truncations tested but did observe a small increase for the reporter with 15 nt 5’ of the UGA codon (Fig. 3B). Furthermore, readthrough efficiency was unaffected by 3’ deletions that were expected to remove the 3’ component of the predicted SL1 stem loop (Fig. 3D). However, reporters with 87 nt or less of *kelch* sequence 3’ of the stop codon had reduced readthrough levels (Fig. 3D). These results suggest that in mammalian cells, SL2 is important for promoting readthrough.

Having identified a sequence supporting *kelch* readthrough in cell culture, we determined whether this sequence promotes readthrough in Drosophila. We modified a dual-luciferase readthrough reporter system so that GFP was fused to the C-terminus of firefly luciferase, allowing readthrough to be monitored using GFP production. As *kelch* readthrough product was highest in the CNS, we assayed readthrough expression from lysates prepared from adult heads in which expression was driven by *nSybGal4*, a strong and specific pan-neuronal Gal4 driver (44). The readthrough efficiency in this reporter system for the 408 bp control fragment (84 nt 5’-UGA-321 nt 3’) was 30% (Fig. 3E, F), significantly higher than we observed in HEK-293T cells, but consistent with the high level of readthrough product we observed when immunoblotting for Kelch in the CNS (Fig. 1A).

The dual-luciferase/GFP reporter constructs used in flies lacked the StopGo sequences that were present in our cell culture assay system (Fig. 3A), raising the possibility that the lack of the StopGo sequences could affect the measured readthrough efficiencies in flies (42). To address this, we made two additional constructs in which the 408 bp UGA readthrough fragment and a UGG IFC were flanked by StopGo (T2A) sequences and inserted between mCherry and GFP (Fig. S2A). Expression of these constructs in neurons using *nSybGal4* produced readily detectable GFP readthrough product (Fig. S2B – E). Fluorescence intensity measurements revealed a readthrough efficiency of 30% (Fig. S2F), equal to that observed when analyzing the same 408 bp fragment in the dual-luciferase/GFP system. These results from two independent experimental approaches indicate that the baseline readthrough efficiency for the 408 bp fragment in Drosophila neurons is 30%.

We next examined the 3’ sequence requirements for readthrough in Drosophila. As was the case in HEK-293T cells, removal of the 3’ half stem of SL1 did not affect the readthrough efficiency (Fig. 3 E, F). In contrast to results in HEK-293T cells, 87 nt 3’ of the stop codon was sufficient to promote high-level readthrough in adult brains. When the sequence was truncated to 51 nt 3’ of the stop codon, completely destroying predicted SL2, we observed a significant decrease in readthrough efficiency (Fig. 3E, F). The basis for the discrepancy between our cultured cell and Drosophila assays for the construct with 87 nt 3’ of the stop codon is not clear although one possibility could be that a Drosophila-specific SL2 binding protein may still interact with the shortened 15 bp stem loop that could still form (Fig. 3C); or perhaps this shortened stem is sufficient to stimulate readthrough in fly but not in human cells. Our analysis demonstrated that at least 51 nt 3’ of the *kelch* UGA are necessary for high-level readthrough and suggest that SL2 contributes to *kelch* readthrough efficiency.

### *kelch* cDNA readthrough reporter highlights cell-specific regulation of readthrough

Our survey of endogenous *kelch* readthrough suggested that the CNS exhibited the highest levels of readthrough, while readthrough product was undetectable in other tissues. To better understand the cell types that support readthrough, we expressed the luciferase/GFP cDNA reporter constructs in distinct cell types. By comparing the pattern of Gal4-driven expression observed with the UGG control (IFC) with the pattern of readthrough of the UGA reporter, we identified cell types that support readthrough to varying extents. When expressed using the ubiquitous *α-Tub84BGal4* driver, GFP from the UGG IFC construct revealed high-level expression in cells of the prothoracic gland (outlined in yellow in Fig. 4A). Within the CNS, *α-Tub84BGal4* drove expression in the optic lobes and in clusters of cells in the central brain (both indicated in Fig. 4A), as well as the ventral nerve cord (VNC), with high-level expression in clusters of midline cells (Fig. 4C, C’) that may represent midline glia (46).

**Fig. 4.**
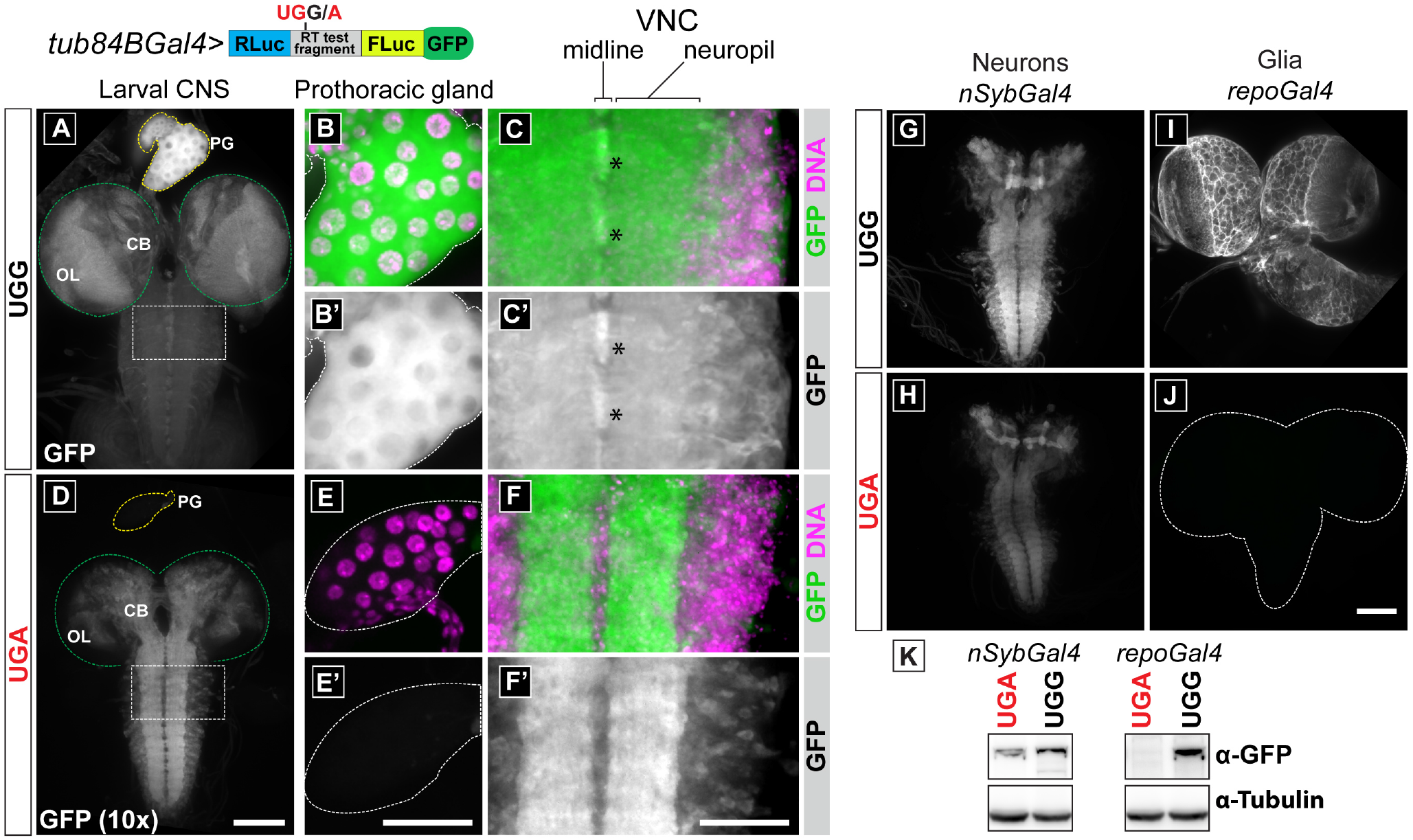
A *kelch* cDNA reporter reveals cell types supporting *kelch* readthrough. (A – C) *α*-Tub84BGa/4-driven expression of the UGG (in-frame control) reporter construct. *α-Tub84BGa/4* drove expression throughout the CNS and at high levels in the endocrine prothoracic gland (PG), outlined in yellow in (A) and shown at higher magnification in B and B’. (C and C’) High magnification view of thoracic neuromeres from the boxed region in (A). High-level GFP expression was observed in clusters of CNS midline cells and throughout cell bodies in the ventral nerve cord (VNC, asterisks). (D – F) Expression of the UGA readthrough reporter driven by *α-Tub84BGal4*. GFP intensity levels of the UGA readthrough reporter were increased 10-fold to allow the pattern of expression to be compared between the UGA and UGG constructs. (D) GFP produced by readthrough accumulated primarily in the central brain and the VNC neuropil. Green dotted line indicates the area occupied by the optic lobes (OL). GFP was not detected in the prothoracic gland (outlined in yellow in (D) and shown at higher magnification in (E and E’)). (F and F’) Higher magnification view of thoracic neuromeres from the boxed region of the VNC in (D). Little GFP was evident in CNS midline cells, and GFP accumulated in fewer cell bodies in the VNC compared to the UGG control. (G-J) Images directly comparing GFP reporter expression in neurons or glia. Expression of reporters specifically in neurons using *nSybGal4* revealed readily detectable readthrough expression from the UGA readthrough reporter (H). (I, J) Expression of reporters specifically in glial cells of the CNS using *repoGal4* failed to detect significant readthrough (region occupied by larval CNS outlined in white in J). (K) Western analysis of reporter expression in lysates prepared from heads of adult flies expressing reporter constructs specifically in neurons or glia. Scale bar for whole larval CNS images (A,D; G-J) = 100 *μ*m. Scale bar for ring gland and VNC high magnification images (B, C, E, F) = 50 *μ*m.

Expression of the UGA readthrough reporter using *α-Tub84BGal4* resulted in an approximate 10-fold reduction in overall fluorescence intensity. To compare the pattern of readthrough (Fig. 4D – F) to the pattern of expression driven by *α-Tub84BGal4* (Fig. 4A – C), the white level for images of the UGA readthrough reporter are increased by 10x. The readthrough reporter produced a markedly different pattern of GFP accumulation compared to the pattern of *α-Tub84BGal4* expression revealed by the IFC. No GFP was observed in cells of the prothoracic gland (Fig. 4E, E’), despite the high level of expression driven in these cells by *α-Tub84BGal4* (Fig. 4A). This demonstrates that some cell types do not support readthrough, even for stop codon contexts that undergo high-level readthrough in other tissues. Within the CNS, GFP resulting from readthrough accumulated predominately in the VNC neuropil as well as in a subset of cell bodies in the central brain region (Fig. 4D) and VNC (Fig. 4D, F and F’). GFP readthrough product did not accumulate in peripheral regions of the optic lobes, where GFP produced from the IFC was present at high levels. Readthrough product was also absent from the clusters of midline cells in the VNC, suggesting that readthrough may not occur in glial cells.

*α-Tub84BGal4* drives expression in many cell types, making it difficult to determine the identity of cells undergoing readthrough in the CNS. To assess the level of *kelch* readthrough supported in neurons, we used *nSybGal4*, a panneuronal driver (Fig. 4G, H). When expressed exclusively in neurons, a readthrough product from the UGA reporter was readily observed (Fig. 4H); fluorescence intensity measurements indicated a readthrough efficiency of approximately 30%. In contrast, when the reporters were expressed using a glial-specific Gal4 driver (Fig. 4I, J), readthrough was essentially undetectable (Fig. 4J). We confirmed these results by immunoblotting extracts from larval brains and observed high levels of readthrough product in neurons but not in glial cells (Fig. 4K). Taken together, this analysis shows that sequences flanking the *kelch* stop codon promote stop codon readthrough in a highly regulated cell-specific manner.

### Highly efficient stop codon readthrough is common in the CNS

A large number of genes are predicted to undergo stop codon readthrough in Drosophila (26–28, 47), including many that are expressed or known to function in the nervous system. Given the high-level readthrough we observed with *kelch* in neuronal tissues, we wondered whether neurons in Drosophila might generally support high-level readthrough. To explore this possibility, we obtained antibodies for genes known or predicted to undergo readthrough for which readthrough would result in a detectable shift in molecular weight. We probed immunoblots of lysates prepared from tissues that supported low-level *kelch* readthrough (ovary) or high-level neuronal *kelch* readthrough (adult brain or larval CNS) (Fig. 5A – D). Western analysis of the RacGEF Sponge (48) revealed an additional product in brain lysate consistent with the predicted molecular weight of the annotated readthrough product (Fig. 5B). In adult brains, the apparent readthrough efficiency was 46%, while readthrough in ovary lysates was undetectable, similar to Kelch (Fig. 5A). We observed a similar result for the transmembrane phosphatase Ptp10D: 56% of Ptp10D protein was detected as readthrough product in brains, with only negligible readthrough detected in ovarian lysates (Fig. 5C). Readthrough of *headcase* has been characterized in detail (15), though it’s relative efficiency in different tissues has not been examined. We did not detect Headcase protein in lysates of adult brains, but we observed a similar level of readthrough (59%) as previously reported in lysates of larval CNS (15). Overall expression of Headcase in ovaries was significantly lower, but we were able to measure an apparent readthrough efficiency of 24% in ovarian lysates. Combined with our study of *kelch* readthrough, these results suggest that in Drosophila, neuronal tissues may be generally permissive for high-level stop codon readthrough.

**Fig. 5.**
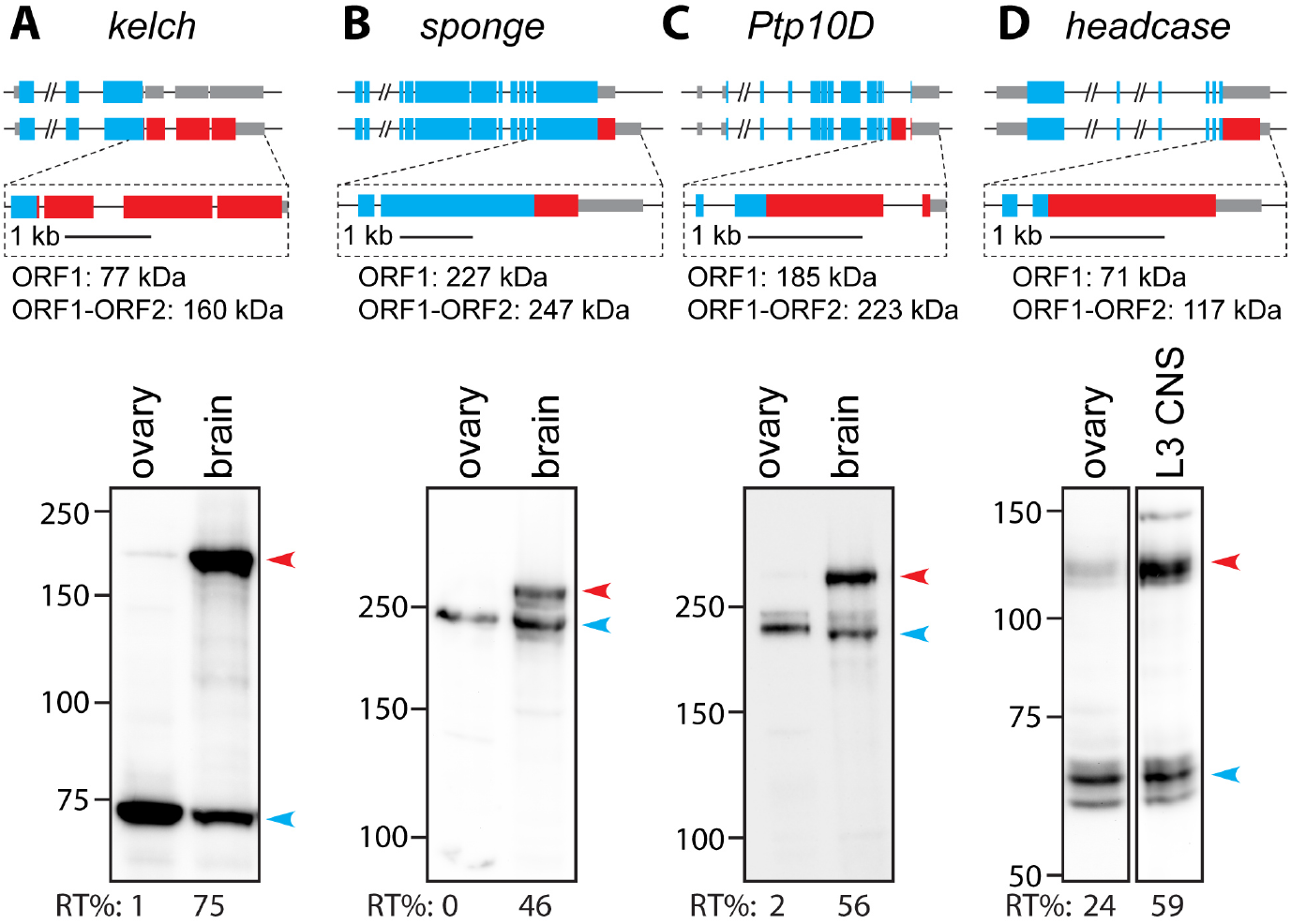
Multiple Drosophila genes show high-level readthrough in neuronal tissue. (A – D) Western blots comparing readthrough translation detected in lysates prepared from ovaries and adult brains (A – C) or third instar larval (L3) CNS (D). Genes encoding each protein are diagrammed at top; blue indicates ORF1 coding sequence, red indicates ORF2 coding sequence. Predicted molecular weights for ORF1 and ORF1-ORF2 readthrough products are listed. Ptp10D is a transmembrane receptor tyrosine phosphatase and is known to be glycosylated, which presumably accounts for the shift in apparent molecular weight (45). Blots were probed with antibodies against the ORF1 product of the indicated genes. Readthrough products are indicated with red arrowheads and ORF1 products are indicated with blue arrowheads. Apparent readthrough efficiencies are indicated beneath each lane, calculated from the ratio of readthrough product to the total quantity of protein detected.

## Discussion

### Highly efficient translational readthrough of Drosophila stop codons

Our results demonstrate that mRNAs from several Drosophila genes undergo high-level, tissue-specific stop codon readthrough in the CNS. For *kelch*, the highest levels of readthrough depend on sequences 3’ of the stop codon. When estimating *kelch* readthrough efficiency based on the steady-state levels of Kelch using an antibody to ORF1, we observed a remarkably high readthrough efficiency of approximately 70% in the CNS. However, differential stability of the ORF1 and readthrough products could significantly bias this estimate. The Kelch ORF1 product is the substrate-binding component of a Cullin-3 RING ubiquitin E3 ligase (CRL3^Kelch^) that can itself be degraded by the ubiquitin proteasome system (36), making it possible that the apparent high-efficiency readthrough was due in part to differential stability of the ORF1 and ORF1-ORF2 products.

To avoid the issue of ORF1 protein instability, we used two independent reporter-based approaches to assess *kelch* readthrough in the CNS. Our readthrough reporter integrated in the *kelch* gene immediately 5’ of the ORF2 stop codon produced a discrete nuclear-localized tdGFP molecule each time readthrough occurred, so that the level of tdGFP served as a reliable indicator of relative readthrough when comparing cells and tissues. This reporter protein was most abundant in imaginal discs and CNS, indicating that the high apparent readthrough efficiency we observed using steadystate measurements was not simply a result of tissue-specific differences in protein stability. In addition, we analyzed *kelch* readthrough using cDNA reporter systems and measured a readthrough efficiency of 30% in the CNS, demonstrating that the CNS supports high-level readthrough of the *kelch* ORF1 stop codon. The large difference in estimated readthrough efficiency observed with the ORF1 antibody immunoblots compared to the cDNA reporter could be due to differential protein stability as discussed above, and/or a lack of sufficient flanking sequence in the reporter constructs necessary for maximal readthrough. Regardless, when taken together with our estimate of similar high-efficiency readthrough in the CNS for *sponge, Ptp10D*, and *headcase*, these results suggest that in Drosophila, the CNS supports high-efficiency developmentally-regulated stop codon readthrough.

### Tissue- and cell type-specific regulation of readthrough

Analysis of the two Drosophila readthrough reporter systems revealed a remarkable degree of tissue- and cell-specific readthrough. Cell-type specific expression of the cDNA reporter revealed highest readthrough efficiency in neurons, while glia and cells of the prothoracic gland terminated translation at the *kelch* ORF1 stop codon with near complete efficiency. We note that *headcase* and *Synapsin*, two Drosophila genes for which readthrough has been characterized, exhibit high efficiency readthrough in the CNS (this study and (15, 49). In addition, genes identified with potential readthrough extensions based on codon conservation were found to be enriched for genes with expression in the nervous system (27), and readthrough expression in the CNS was verified for at least one of these, *Abd-B* (26). Interestingly, an analysis of premature stop codon (PTC) readthrough in Drosophila revealed highly efficient readthrough of PTCs in CNS neurons, but not glia (50).

Recent studies indicate that regulated translational readthrough in cellular gene decoding may be more widespread than previously appreciated. Cell-type specific regulation of readthrough was observed in mice using ribosome profiling (51), suggesting broader translational mechanisms that permit cell-type specific readthrough in flies and mammals. Human AQP4 encodes an aquaporin that undergoes efficient stop codon readthrough in cell culture (13) and for which readthrough efficiency may be regulated in a tissuespecific manner (52).

### Possible mechanisms controlling readthrough

A number of factors are known to influence the efficiency of stop codon readthrough including the local sequence context as well as the formation of RNA secondary structures 3’ of the stop codon. In addition, trans-acting factors have been shown to influence readthrough efficiency, including release factor abundance or modification, tRNA abundance, as well as other protein factors (53, 54).

Our analysis of *kelch* suggests that a predicted mRNA stem-loop 3’ of the stop codon stimulates high-efficiency stop codon readthrough. Evidence for a stem loop structure promoting readthrough has also been described for *head-case* (15), and a conserved sequence 3’ of the *Ptp10D* stop codon is also predicted to form a stem loop (28), Fig. S3). Similar to *kelch*, readthrough of the *headcase* and *Ptp10D* stop codons appears to be developmentally regulated so that high-efficiency readthrough of these genes is promoted in the CNS, suggesting that stem loop structures may be important for promoting high-efficiency stop codon readthrough in the CNS.

One possible role of stem loops or other RNA secondary structures in enhancing readthrough could be precise pausing of ribosomes when a stop codon is within the A site, thus increasing dwell time and potentially nudging competition in favor of aminoacyl-tRNA decoding. Recently, cryo-EM studies revealed that during stop codon recognition by eukaryotic release factor 1 (eRF1), the nucleotide immediately 3’ of the stop codon is pulled into the A site to form a compact U-turn (2, 3). Perhaps a stable RNA secondary structure at the mRNA entrance tunnel could prevent eRF1 from pulling the 4th nucleotide into the A site and thus reduce termination efficiency. Another possibility is that a trans-acting factor could stimulate readthrough by binding to and stabilizing the RNA structure. Although there are no known examples of this type used for readthrough, recently two examples of viral ribosomal frameshifting signals exist where frameshifting is trans-activated through the action of viral (cardiovirus 2A, arterivirus nsp1*β*) and cellular proteins (poly(C) binding protein) (55, 56).

### Evasion of NMD does not explain tissue-specific readthrough

A consideration with stop codons that are read through, especially when the stop codon is followed by a second long ORF, is nonsense-mediated decay (NMD). Indeed, avoidance of NMD has been documented in a few instances of readthrough in RNA viruses (18, 57). Proximity of the stop codon to a 3’ splice junction, and so to exon junction complex (EJC) proteins on the mRNA is relevant in mammals where any stop greater than 50 nt upstream of an EJC is marked for NMD (58). However, it is unlikely that differential tissue-specific resistance to NMD due to proximity to an EJC could explain our observations since the *Ptp10D* stop codon is 1,045 nt from a 3’ splice site. In addition, splice junctions and EJC proteins do not appear to be involved with PTC recognition for NMD in Drosophila (59, 60) and an instance of PTC suppression found to occur in the CNS is not due to tissue-specific differences in the NMD response (50). Furthermore, publicly available Drosophila RNAseq data available at FlyBase and FlyAtlas2 reveal comparable transcript levels in low-readthrough (ovary, salivary gland) versus high-readthrough (brain) tissues, for both *kelch* and the other readthrough genes (61, 62). Thus, it is unlikely that our results can be explained by a lower level of NMD in neurons.

Recently, it has become clear that the position of a stop codon within the mRNA can affect termination efficiency (63) with stop codons distant from the polyA tail having lower termination efficiency, typically resulting in NMD (58). Intriguingly, a novel genetic code was recently discovered where all three standard stop codons (TAA, TAG, and TGA) specify amino acids in *Condylostoma magnum*. These codons are decoded as amino acids at internal positions, but specify translation termination when in close proximity to an mRNA 3’ end (64, 65). However, the length of the 3’ UTR after the first stop codon in Drosophila readthrough genes ranges widely from a few nt to thousands (26, 31). For example, *kelch* mRNA is abundant in ovarian cells where readthrough is nearly undetectable even though the 3’ UTR following the first stop codon is over 3000 nt. This constellation of readthrough genes in the Drosophila genome argues against a correlation between 3’ UTR length and evasion of NMD in favor of readthrough.

### ORF2 function

Evidence for extensive stop codon readthrough in Drosophila comes from ribosome footprinting experiments (31) and analysis of codon conservation downstream of the first stop codon in genes (26–28, 47). Based on these data, there are now over 400 genes annotated in FlyBase as readthrough genes with highly conserved extensions among Drosophila species. In contrast, there are only a few dozen readthrough genes documented in the mouse/human genome (13, 31, 51, 66, 67) suggesting the Drosophila genome finds readthrough particularly useful. Curiously, the peptide extensions produced by readthrough genes in Drosophila vary considerably in length from fewer than 10 amino acids to over 800. Remarkably, many of these readthrough extensions have been conserved over hundreds of millions of years of insect evolution (28), indicating that they are under positive selection and function to increase the fitness of these insects. That readthrough is exceptionally and dramatically efficient in the central nervous system is intriguing.

## Materials and Methods

### Cloning and molecular biology

For the *kelch* readthrough reporter, we designed a homology donor (HD) construct with a readthrough reporter cassette located immediately 5’ of the *kelch* ORF2 stop codon. The reporter cassette consisted of a 3xFLAG epitope tag, a T2A StopGo sequence and a nuclear-localized tandem green fluorescent protein (NLS::tdGFP) reporter gene. 5’ and 3’ homology arms were 1,501 bp and 1,100 bp, respectively. A 3xP3-DsRed transformation marker flanked by PiggyBAC transposon termini (derived from pScarlessHD-DsRed; gift of Kate O’Connor-Giles, Addgene plasmid #64703) was inserted between duplicated TTAA PiggyBAC target sequences in the 3’ homology arm, 42 nt 3’ of the ORF2 stop codon. A 1.5 kb fragment containing the readthrough reporter cassette and adjacent sequences was synthesized (Genscript) and assembled with remaining 5’ and 3’ homology sequences and the DsRed transformation marker using conventional cloning techniques. The protospacer adjacent motif (PAM) of the guide RNA (gRNA) site used for targeting was mutated from AGG to AGA in this vector. Oligos encoding a gRNA targeting a site 40 nucleotides 5’ of the ORF2 TAA stop codon (gRNA: gGCTGTTGGTAGTGCTT|GGA; | indicates cleavage site. Initial G residue not present in genomic sequence) were cloned into *pBFvU6.2* (68). The homology donor construct and U6-gRNA plasmids were injected into fly embryos expressing transgenic Cas9 at BestGene (Houston TX). *DsRed^+^* transformants were isolated, and the *DsRed^+^* marker gene was subsequently excised by crossing to PiggyBac transposase. Initial HDR events and DsRed-excision lines were verified by PCR for correct targeting and integration. For the generation of dual luciferase expression constructs for mammalian cell expression, inserts were amplified by PCR using a *kelch* synthetic DNA (gBlock – Integrated DNA Technologies: IDT) as template and cloned into pSGDluc (42) using standard techniques. For Drosophila constructs, readthrough test fragments flanked by dual luciferase coding sequences were cloned into a modified pJFRC28, a PhiC31 UAS-GFP expression vector (69). Plasmids were integrated at the *attP2* site on chromosome 3L by injection at Rainbow Transgenics (Camarillo, CA) or Genetivision (Houston, TX). All constructs were verified by DNA sequencing.

### Immunoblotting

For blots analyzing readthrough in different tissues (Fig. 1), samples were dissected in physiological buffers (PBS, IMADS (70), or Schneider’s S2 medium (ThermoFisher), flash frozen on dry ice and stored at −80°C until use. Ovarian tissue was lysed in SDS sample buffer using plastic pestles and a motorized homogenizer (Kontes). All other tissues were lysed by freeze-thaw treatment, pipetting and incubation at 95°C in SDS sample buffer. Quantities of tissue used for equal loading were determined empirically; equal loading was verified by amido black staining. Proteins were separated on either a NuPAGE 4 – 12% gradient gel (Fig. 1C) or 7.5% polyacrylamide gels (all other figures). Separated proteins were transferred to nitrocellulose and membranes were blocked and probed with the following antibodies: mouse *α*-Kelch (Kel 1B, DSHB, 1:25), mouse *α*-Adducin (1B1, DSHB, 1:50), mouse *α*-FLAG (M2, Sigma F1804, 1:1,000), rabbit *α*-GFP (Invitrogen A-11122, 1:1,000), and mouse *α*-actin (JLA20, DSHB, 1:50). Blots were washed, probed with corresponding secondary HRP-conjugated secondary antibodies (Pierce), washed again, incubated with ECL reagents (Clarity, BioRad) and imaged using a CCD (ProteinTech).

For semi-quantitative immunoblotting of GFP readthrough products, lysates were prepared from heads expressing UAS reporter constructs. 10 heads per biological replicate were flash frozen on dry ice, homogenized in 50 *μl* of SDS sample buffer, and incubated at 95°C for 5 minutes. 10 *μl* were loaded per lane, separated on 7.5% polyacrylamide gels and blotted as described above. Intensities of GFP and Adducin bands were measured in FIJI, and GFP was normalized for loading based on Adducin levels. Readthrough efficiency was calculated as the fraction of GFP detected in the UGA readthrough sample relative to its matched UGG in-frame control.

### Immunofluorescence

Adult brains were dissected in Schneider’s S2 medium, fixed in 2% paraformaldehyde (PFA) for 55 min and washed 4x in PBTx (PBS + 0.1% Triton X-100, 0.5 % BSA). All other tissues were dissected in PBS, fixed in a 4% PFA with or without 0.1% Triton X-100 and washed 4x in PBTx. Larval CNS expressing GFP and mCherry from cDNA reporter constructs were incubated with 0.5 *μ*g/ml 4,6-diamino-2-phenylindole (DAPI), washed, and mounted in 90% glycerol/20 mM Tris pH 8.0. Samples from the NLS::tdGFP *kelch* readthrough reporter were blocked for 2 hours in PBTx + 5% normal goat serum (NGS) and incubated with the following primary antibodies overnight in PBTx + 5% NGS: mouse anti-lamin (ADL67.10, DSHB, 1:7) and rabbit anti-GFP (ThermoFisher A11122, 1:500). Adult brains were incubated for two overnights with *α*-Rabbit 488 (ThermoFisher A11034, 1:500) and 0.5 *μ*g/ml DAPI, washed and mounted in ProLong (ThermoFisher). Other tissues were washed 4x in PBTx and incubated overnight with goat *α*-Rabbit 488 (1:500) and goat *α*-mouse 568 (Invitrogen A11031, 1:500). These tissues were washed and mounted in ProLong.

### Imaging and analysis

Samples were imaged using either a Leica SP8 scanning confocal system using a 40x Plan Apo 1.30 NA objective or a Zeiss Axio observer 7 inverted microscope with a CrEST X-light v2 spinning disc system, Photometrics BSi sCMOS camera and either a 20x Plan Apo 0.8 NA objective or a 40x C-Apo 1.2 NA water-immersion objective. Images were processed and analyzed using ImageJ/FIJI. For image sets in which fluorescence intensities were compared, images were collected under identical imaging conditions using the Zeiss/sCMOS system. Images were processed in FIJI and levels were set identically except where indicated in the figure legend. For quantification of readthrough of the mCherry/GFP readthrough reporter, summed-intensity projection images of larval brains expressing mCherry and GFP were analyzed. Masks outlining each CNS were generated by auto-thresholding the mCherry channel of each image in FIJI (n = 4 for each construct), and GFP intensities within the masked region were measured. Readthrough was calculated based as the ratio of the summed intensity of GFP in the TGA readthrough construct to the TGG in-frame control.

### Cell Culture and Transfections

HEK293T cells (ATCC) were maintained in DMEM supplemented with 10% FBS, 1 mM L-glutamine and antibiotics. Cells were transfected with Lipofectamine 2000 reagent (Invitrogen), using the 1-day protocol in which suspended cells are added directly to the DNA complexes in half-area 96-well plates. The following were added to each well: 25 ng of each plasmid plus 0.2 *μ*l Lipofectamine 2000 in 25 *μ*l Opti-Mem (Gibco). The transfecting DNA complexes in each well were incubated with 3 × 104 cells suspended in 50 *μ*l DMEM + 10% FBS at 37°C in 5% CO_2_ for 24 hr.

### Dual Luciferase Assay

Firefly and Renilla luciferase activities were determined using the Dual Luciferase Stop & Glo Reporter Assay System (Promega). Relative light units were measured on a Veritas Microplate Luminometer with two injectors (Turner Biosystems). Transfected cells were lysed in 12.6 *μ*l of 1 × passive lysis buffer (PLB) and light emission was measured following injection of 25 *μ*l of either Renilla or firefly luciferase substrate. Readthrough efficiencies (% readthrough) were determined by calculating relative luciferase activities (firefly/Renilla) of TGA constructs and dividing by relative luciferase activities from replicate wells of control TGG constructs. The number of biological replicates for each experiment is indicated in each figure legend.

## ACKNOWLEDGEMENTS

We thank Jenna O’Neil for preliminary work on this project at Yale. N.W. and J.A. thank Ray Gesteland for facilitating the Utah component of this work. This research was supported by Irish Research Council Advanced Laureate (IRCLA/2019/74) to J.F.A and NIH grant R01 GM0043301 to L.C.

## Supplementary Information

**Fig. S 1.**
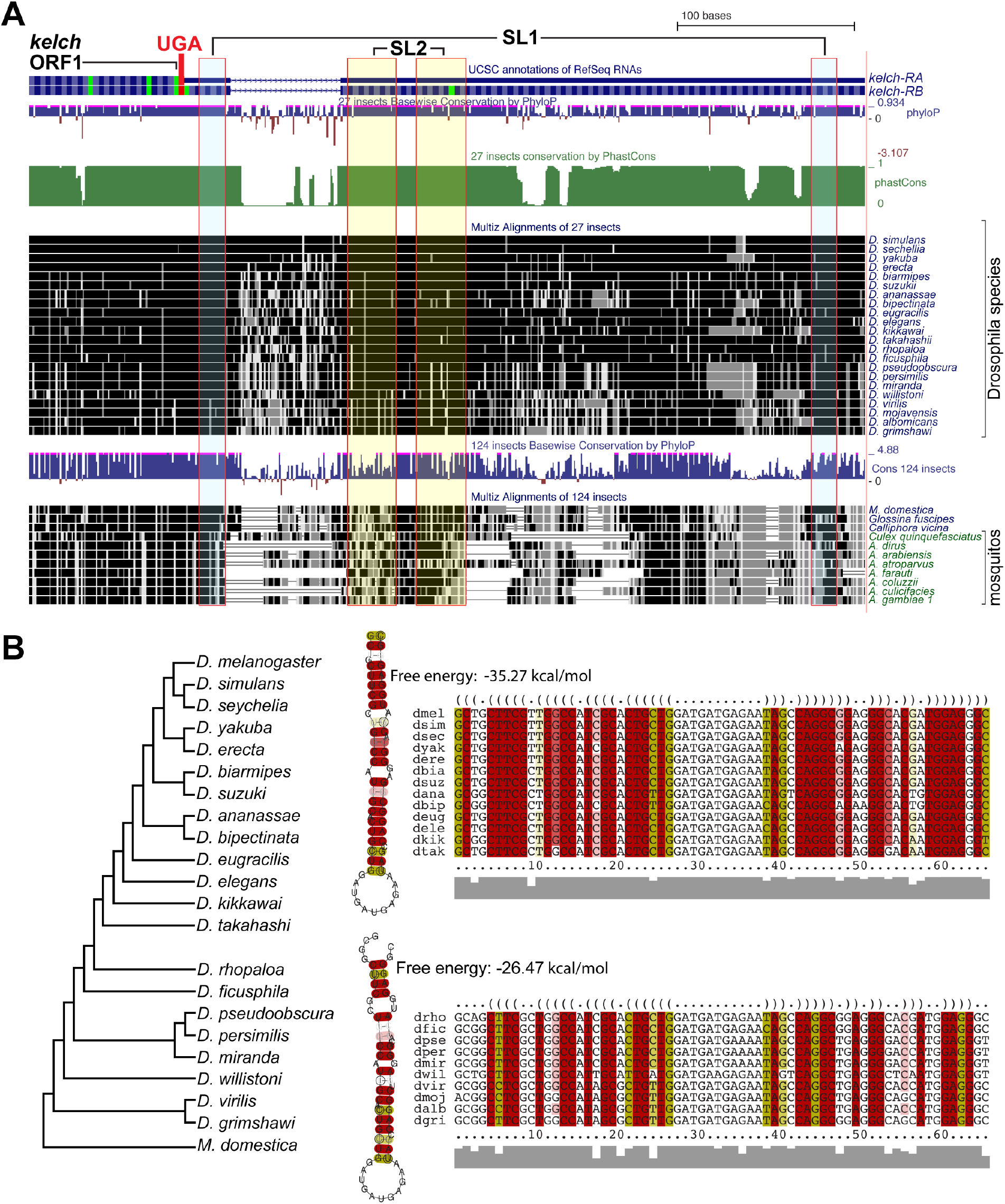
Conservation of predicted stem-loop sequences 3’ of the *kelch* UGA stop codon. A. Screenshot of UCSC genome browser showing conservation of nucleotide sequences involved in stem-loop pairing (SL1 highlighted in cyan, SL2 in yellow). B. Conservation of pairing sequences in RNA stem loop SL2 based on RNAalifold conservation-based structure prediction (1). Species closely related to *D. melanagaster* show potential to form stem loop (top alignment). More distantly related species retain potential to form a similar stem loop.

**Fig. S 2.**
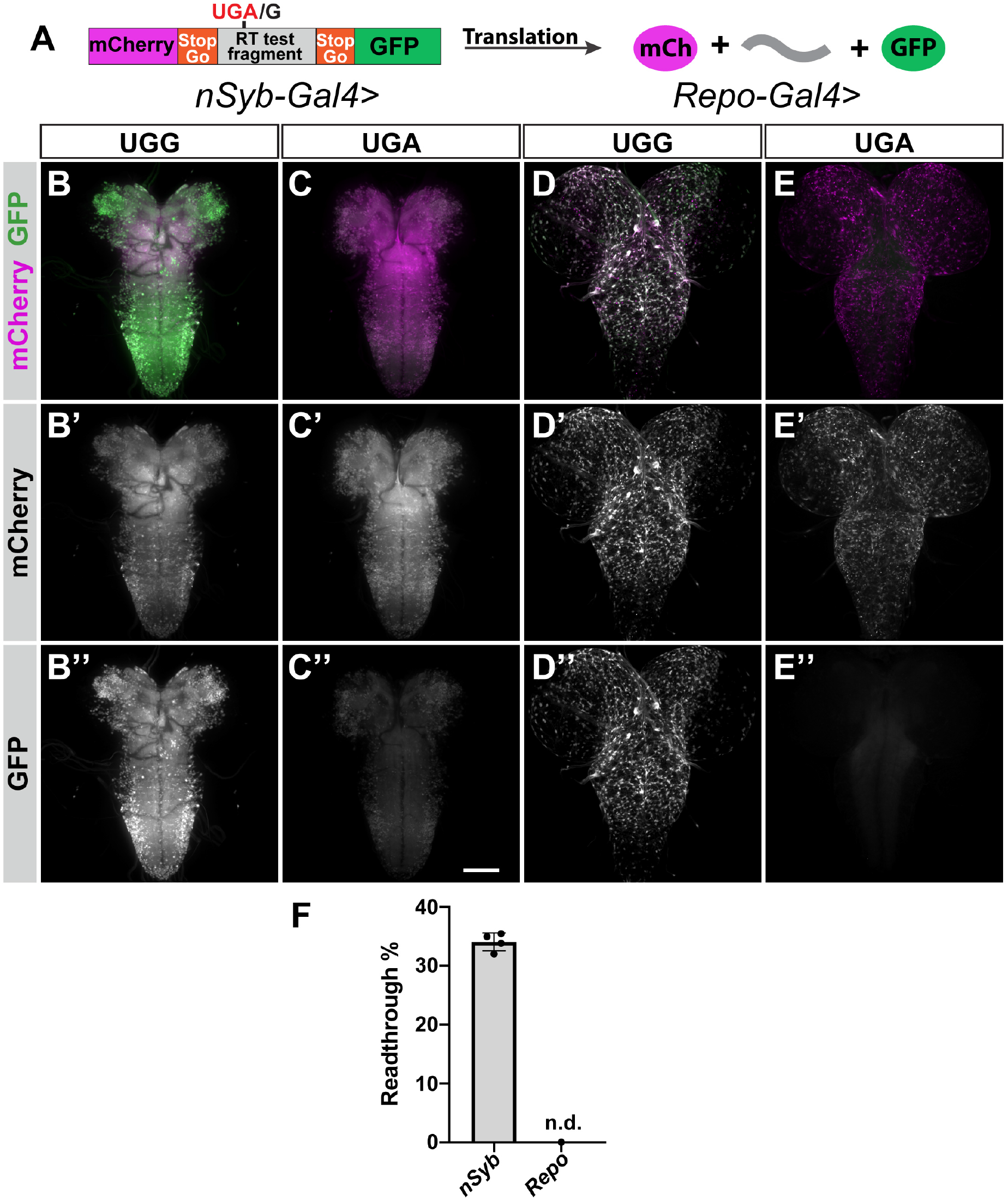
Dual-fluorescent readthrough reporter. A. Diagram of dual-fluorescent readthrough reporter. T2A StopGo sequences flank readthrough test fragment, resulting in production of mCherry and GFP without associated fusion sequence. When expressed in neurons, GFP expression produced from UGA construct (C”) reveals 30% readthrough efficiency relative to UGG in-frame control (B). Quantification based on summed fluorescence intensities of neuronal GFFP expression is plotted in F. When expressed in glia, GFP produced by the UGA readthrough reporter was not be detected above background.

**Fig. S 3.**
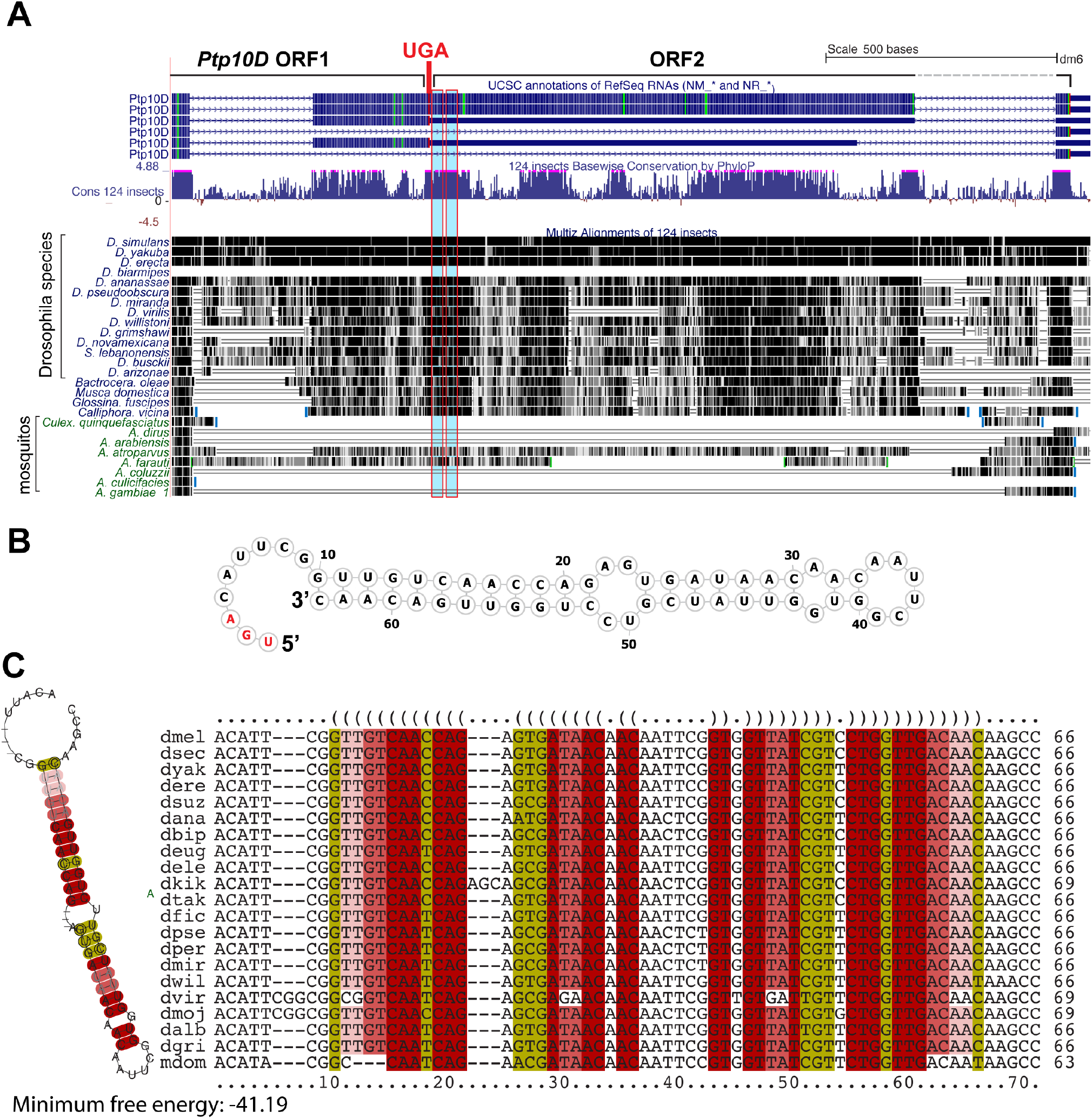
Conservation of predicted stem-loop sequences 3’ of the *Ptp10D* UGA stop codon. A. Screenshot of UCSC genome browser showing nucleotide conservation of sequences involved in stem-loop pairing (sequences encoding 5’ and 3’ stems are highlighted in cyan). B. Stem loop predicted by RNAfold from the Vienna Package (2). ORF1 UGA termination codon is marked in red. C. RNAal-ifold structure prediction based alignment of related species (1). Conservation of predicted pairing sequences is present throughout Drosophila.

## Notes

### Competing Interest Statement

The authors have declared no competing interest.

